# *In Silico* Identification and Experimental Validation of Novel KPC-2 β-lactamase Inhibitors

**DOI:** 10.1101/396283

**Authors:** R. Klein, P. Linciano, G. Celenza, P. Bellio, S. Papaioannou, J. Blazquez, L. Cendron, R. Brenk, D. Tondi

## Abstract

Bacterial resistance has become a worldwide concern, particularly after the emergence of resistant strains overproducing carbapenemases. Among these, the KPC-2 carbapenemase represents a significant clinical challenge, being characterized by a broad substrate spectrum that includes aminothiazoleoxime and cephalosporins such as cefotaxime. Moreover, strains harboring KPC-type β-lactamases are often reported as resistant to available β-lactamase inhibitors (clavulanic acid, tazobactam and sulbactam). Therefore, the identification of novel non β-lactam KPC-2 inhibitors is strongly necessary to maintain treatment options. This study explored novel, non-covalent inhibitors active against KPC-2, as putative hit candidates. We performed a structure-based *in silico* screening of commercially available compounds for non-β-lactam KPC-2 inhibitors. Thirty-two commercially available high-scoring, fragment-like hits were selected for *in vitro* validation and their activity and mechanism of action *vs* the target was experimentally evaluated using recombinant KPC-2. N-(3-(1H-tetrazol-5-yl)phenyl)-3-fluorobenzamide (**11a**), in light of its ligand efficiency (LE = 0.28 kcal/mol/non-hydrogen atom) and chemistry, was selected as hit to be directed to chemical optimization to improve potency *vs* the enzyme and explore structural requirement for inhibition in KPC-2 binding site. Further, the compounds were evaluated against clinical strains overexpressing KPC-2 and the most promising compound reduced the MIC of the β-lactam antibiotic meropenem by four fold.

## Introduction

The emergence of KPC-2 class-A Beta-Lactamase (BL) carbapenemase, which confers resistance to last resort carbapenems, poses a serious health threat to the public. KPC-2, a class A BL, uses a catalytic serine to hydrolyze the β-lactam ring. Specifically, the hydrolysis reaction proceeds through a series of steps involving: (i) the formation of a precovalent complex, (ii) the conversion to a high-energy tetrahedral acylation intermediate, (iii) followed by a low-energy acyl-enzyme complex, (iv) a high-energy tetrahedral de-acylation intermediate consequent to catalytic water attack, and (v) finally the release of the hydrolyzed β-lactam ring product from the enzyme. [1–6].

Notably to treat infections caused by bacteria that produce class A BLs, mechanism-based inhibitors (i.e., clavulanic acid, sulbactam, and tazobactam) are administered in combination with β-lactam antibiotics. However, strains harboring KPC-type β-lactamases are reported to be resistant to available β-lactamase inhibitors. Moreover, because of KPC-2’s broad spectrum of activity (which includes penicillins, cephalosporins, and carbapenems) treatment options against KPC-2-producing bacteria are scarce, and “last-resort” carbapenems are ineffective as well [7]. Therefore, studies directed to the discovery of novel, non β-lactam KPC-2 inhibitors have multiplied in the last years. Recently, new drugs able to restore susceptibility to β-lactams i.e. the novel inhibitor avibactam in combination with ceftazidime (CAZ) and RPX7009 (vaborbactam) with meropenem have been approved (Fig. 1)[8-10].

**Figure 1.**
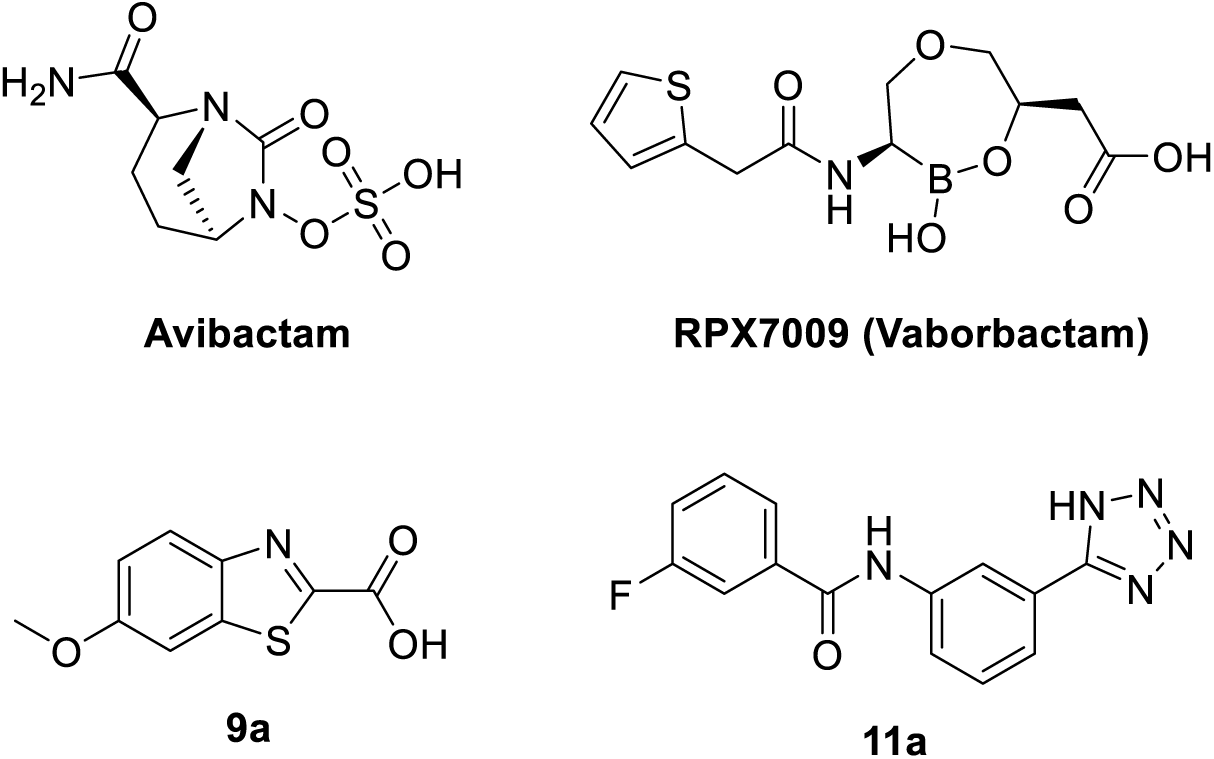
Chemical structure of avibactam, RPX7009, and compounds **9a** and **11a**

As attention on KPC-2 rises, the number of crystal structures of its apo and complexed form disclosed in the PDB has increased, making KPC-2 a druggable target for structure based drug design efforts and for the study of novel, non β-lactam like inhibitors of this threatening carbapenemase [9–12]

Recently, two crystal structures of the hydrolyzed β-lactam antibiotics cefotaxime and faropenem in complex with KPC-2 were determined (PDB codes 5UJ3, 5UJ4; Fig. 2).[13]

**Figure 2.**
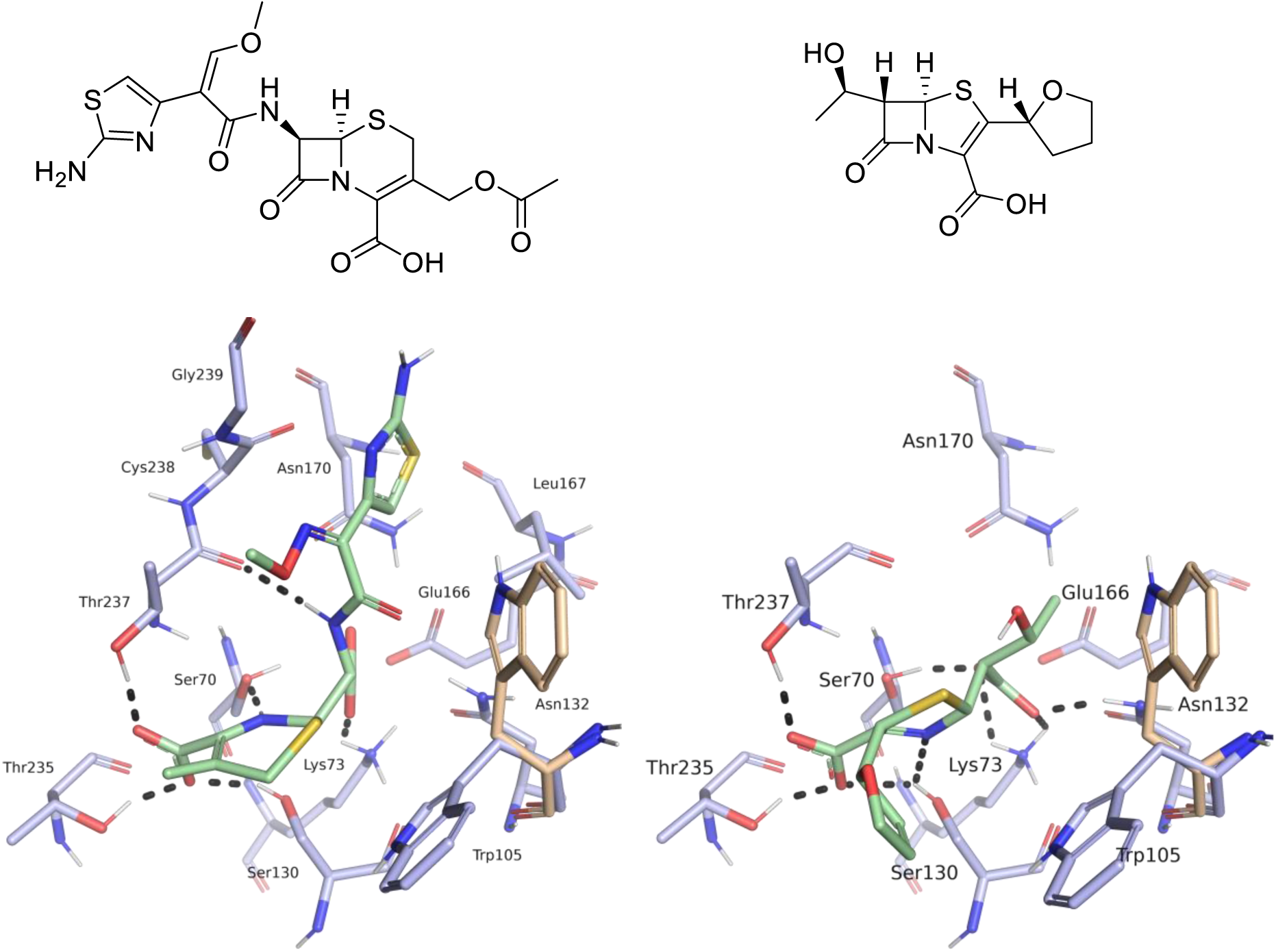
Structures and binding modes of hydrolyzed β-lactam antibiotics in the KPC-2 binding site. Left: binding mode of hydrolyzed cefotaxime (PDB code 5UJ3). Right: binding mode of hydrolyzed faropenem (PDB code 5UJ4). The second rotamer of Trp105 adopted in the apo-enzyme is coloured in beige, protein side chains in blue and ligands in green. Hydrogen bonds are indicated as black dots.

Both ligands form hydrogen-bond interactions with their C4-carboxyl group to Ser130, Thr235 and Thr237. The dihydrothiazine moiety of cefotaxime and the dihydrothiazole moiety of faropenem forms π-π-stacking interactions with Trp105. In the apo-enzyme, this side chain adopts two rotamers, upon binding of a ligand just one. Mutagenesis studies have shown the importance of Trp105 in substrate recognition [7]. The faropenem ring nitrogen forms a hydrogen-bond interaction with Ser130, whereas the ring nitrogen of cefotaxime a hydrogen bond with Ser70. The aminothiazole ring of cefotaxime forms van-der-Waals contacts with Leu167, Asn170, Cys238 and Gly239, while the oxyimino group and the hydroxyethyl group of faropenem are solvent exposed (Fig. 2).[13]

Based on this and other structural information, we used a hierarchical screening cascade for the discovery of non β-lactam like KPC-2 inhibitors. The selected candidates were then validated as hits against isolated recombinant KPC-2. Among the tested compounds **9a**, a benzotiazole derivative, and **11a**, a tetrazole-containing inhibitor, showed the highest activity against KPC-2 and behaved as competitive inhibitors of the targeted carbapenemase (Fig. 1). Compound **11a** was subsequently directed to chemical optimization to improve potency *vs* the enzyme and explore structural requirement for inhibition in KPC-2 binding site. Further, the compounds were evaluated against clinical strains overexpressing KPC-2 and the most promising compound reduced the MIC of the β-lactam antibiotic meropenem by four fold.

## Materials and Methods

### Pharmacophore hypothesis

A search for similar binding sites of KPC-2 was carried out using the online tool PoSSuM - Search K [15,16]. Based on shared ligand interactions in the retrieved structures (Table 1), a pharmacophore was defined based on a *K. pneumoniae* KPC-2 protein structure (PDB code 3RXW) [17] and the ligand OJ6 of CTX-M-9 β-lactamase (PDB code 4DE1) [18]. The derived pharmacophore contained a hydrogen-bond acceptor feature for interaction with Thr237, Thr235 and Ser130, a hydrophobic feature for π-stacking with Trp105 and a hydrogen bond acceptor feature for interactions with Asn132 (Fig. 3).

**Table 1:**
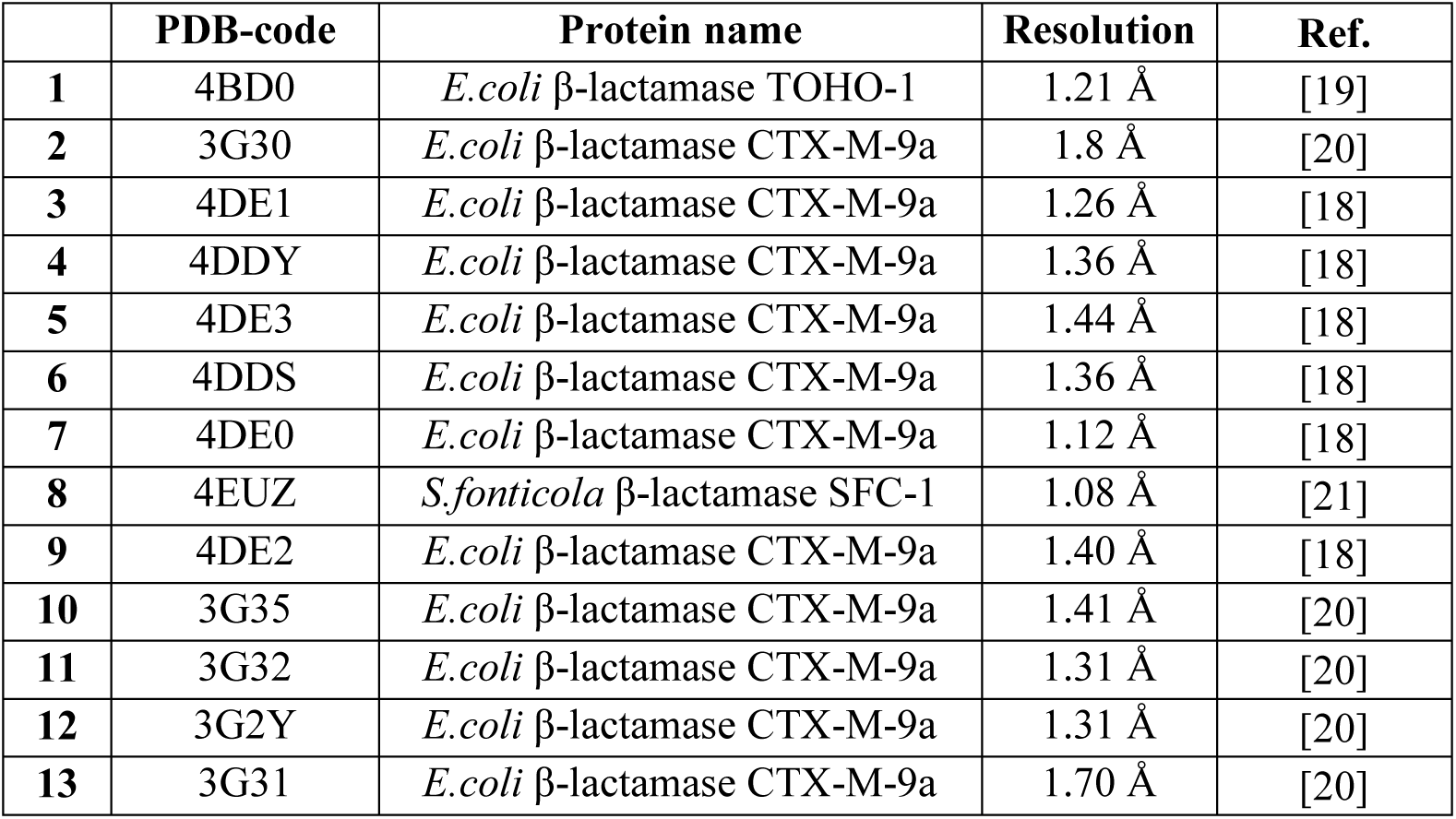
Result of PoSSuM Search K for similar binding sites. Structures with binding sites similar with structure 3RXW in complex with a non-covalent ligand were reported.

|  | PDB-code | Protein name | Resolution | Ref. |
| --- | --- | --- | --- | --- |
| 1 | 4BD0 | *E.coli* β-lactamase TOHO-1 | 1.21 Å | [19] |
| 2 | 3G30 | *E.coli* β-lactamase CTX-M-9a | 1.8 Å | [20] |
| 3 | 4DE1 | *E.coli* β-lactamase CTX-M-9a | 1.26 Å | [18] |
| 4 | 4DDY | *E.coli* β-lactamase CTX-M-9a | 1.36 Å | [18] |
| 5 | 4DE3 | *E.coli* β-lactamase CTX-M-9a | 1.44 Å | [18] |
| 6 | 4DDS | *E.coli* β-lactamase CTX-M-9a | 1.36 Å | [18] |
| 7 | 4DE0 | *E.coli* β-lactamase CTX-M-9a | 1.12 Å | [18] |
| 8 | 4EUZ | *S.fonticola* β-lactamase SFC-1 | 1.08 Å | [21] |
| 9 | 4DE2 | *E.coli* β-lactamase CTX-M-9a | 1.40 Å | [18] |
| 10 | 3G35 | *E.coli* β-lactamase CTX-M-9a | 1.41 Å | [20] |
| 11 | 3G32 | *E.coli* β-lactamase CTX-M-9a | 1.31 Å | [20] |
| 12 | 3G2Y | *E.coli* β-lactamase CTX-M-9a | 1.31 Å | [20] |
| 13 | 3G31 | *E.coli* β-lactamase CTX-M-9a | 1.70 Å | [20] |

**Figure 3.**
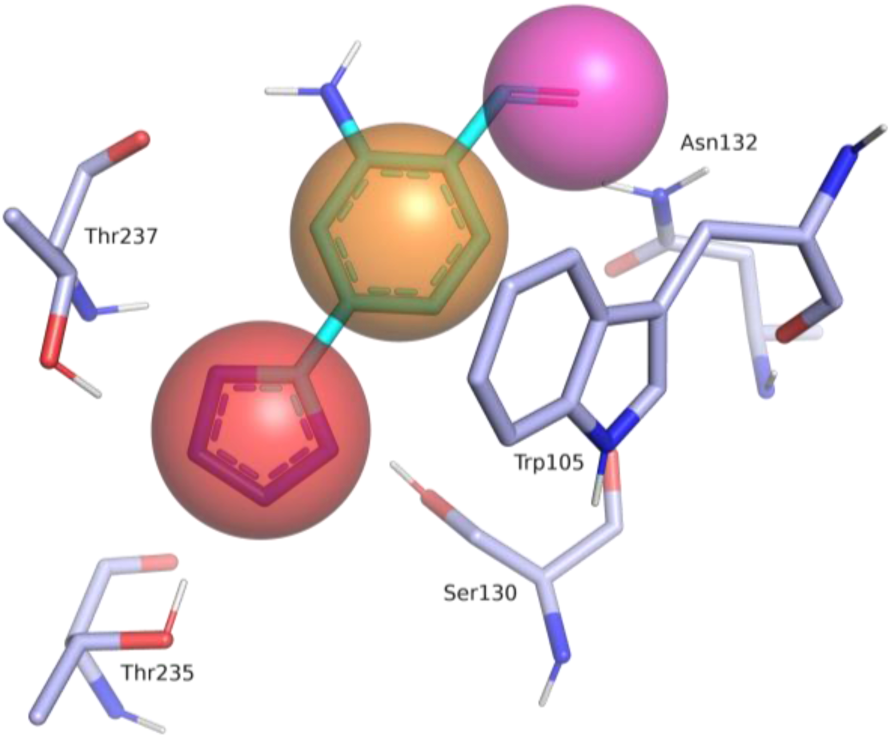
Binding site of KPC2 (PDB code 3RXW, blue) superimposed with a fragment of the ligand OJ6 bound to CTX-M-9 β-lactamase (PDB code 4DE1, cyan) and pharmacophore features (red: hydrogen-bond acceptor, orange: hydrophobic interaction feature, purple: hydrogen-bond donor) (Ambler numbering) [22].

## Virtual Screening

Our in-house MySQL-database of commercially available compounds was filtered for compounds fulfilling the following lead-like criteria: between 10 ten and 25 twenty-five heavy atoms, between one and six hydrogen-bond acceptors, between one and three hydrogen-bond donors and a clog P between - 3 and 3. In addition, the complexity was limited by only including compounds with less than 7 rotatable bonds and between 1 and 3 ring systems. Compounds containing unwanted reactive or toxic functional groups were excluded as well [23].

In-house python scripts based on OpenEye’s OEChem toolkit (OEChem, version 2016.6.1, OpenEye Scientific Software, Inc., Santa Fe, NM, USA) were used to charge, tautomerize and stereoisomerize the selected compounds. Conformers were generated using OpenEye’s OMEGA toolkit [24]. The pharmacophore filtering was carried out using Molecular Operating Environment (MOE, Chemical Computing Group). Compounds that passed the pharmacophore filter were transformed into a format suitable for docking as described previously [25].

The crystal structure of *K. pneumoniae* KPC-2 (PDB code 3RXW) [17] was used as receptor for docking. The ‘protonate 3D’ tool of MOE was used to add polar hydrogen atoms to the receptor, energy minimize their positions and to assign partial charges based on the AMBER force field parameters. Water molecules and ligands (CIT and SR3) were deleted and the position of the Ser69 side chain was energy minimized with the same force field parameters. The structure was aligned with the crystal structure of *E.coli* CTX-M-9 (PDB code 4DE1) and the ligand 0J6 was used to define spheres as matching points for docking. Grid-based excluded volume, van-der-Waals potential and electrostatic potential as well as solvent occlusion maps were calculated as described earlier [26,27].

The compounds were docked into the binding site of KPC-2 using DOCK3.6 [27–29]. Parameters for sampling ligand orientations were set as follows: bin size of ligand and receptor were set to 0.4 Å, overlap bins were set to 0.2 Å and the distance tolerance for receptor and ligand matching spheres was set to 1.5 Å. Each docking pose which did not overlap with the receptor was scored for electrostatic and van-der-Waals complementarity and penalized according to its estimated partial desolvation energy. For each compound, only the best-scoring pose out of its tautomers, protonation states or ring alignments was saved in the final docking hit list. The docking hit list was filtered with the pharmacophore described above, keeping the ligand positions rigid. Compounds passing this filter were ranked by their calculated ligand efficiency [30,31] and inspected by eye.

## Expression and purification of recombinant KPC-2

The *bla*_KPC-2_ gene was kindly provided by Prof. Sergei Vakulenko (University of Notre Dame du Lac, Indiana, USA) and cloned as already reported [32] and transformed into competent *E.coli* BL21 (DE3) cells for protein expression. 50 mL of Tryptic Soy Broth (TSB) (50 mg/L kanamycin) were inoculated with fresh colonies and grown at 37°C. 4 mL of the overnight culture was used to inoculate 1.3 L of TSB (50 mg/L kanamycin) grown at 37°C with shaking to an optical density of 0.5 measured at 600 nm. Then expression of recombinant *bla* gene was induced by adding 1.0 mM IPTG (isopropyl-D-thiogalactopyranoside) and the cells were again allowed to grow at 20 °C overnight. Bacteria were harvested by centrifugation at 4000 rpm for 20 minutes. The pelleted cells were resuspended in Tris-HCl 50 mM pH 7.4-7.5. Periplasmatic proteins were extracted as reported in the pET System Manual (TB055 10th Edition Rev. B 0403) and subsequently dialyzed in sodium acetate buffer (50 mM, pH 5.0). The protein was conveniently purified in a single step using a Macro-Prep High S resin and eluted using sodium acetate 50 mM pH 5.0 and a sodium chloride (NaCl) linear gradient from 100 to 500 mM. The purified protein was dialyzed overnight in sodium phosphate buffer 50 mM, pH 7.0 [32,33].

## Inhibition Assays

The hydrolytic activity of KPC-2 activity was measured using the β-lactam substrates CENTA (100 uM, K_M_ 70 uM) or nitrocefin (114 uM, KM 36 μM) in reaction buffer consisting of 50 mM of PB at pH 7.0 at 25°C with 0.01% v/v Triton X-100 to avoid compound aggregation and promiscuous inhibition.[34] Reactions were monitored using a Beckmann DU640^®^ spectrophotometer at 405nM for CENTA and 480 nM wavelength for nitrocefin [35]. The test compounds were synthesized as described below or purchased from Enamine, TimTec, Vitas-M, ChemBridge, Otava, Life Chemicals or Apollo Scientific and assayed without further purification. Compounds were dissolved in dimethyl sulfoxide (DMSO) to a concentration of 25 mM and stored at −20°C. The highest concentration at which the compounds were tested was up to 1 mM (depending on their solubility). All experiments were performed in duplicate and the error never exceeded 5%. The reaction was typically initiated by adding KPC-2 to the reaction buffer last. To control for incubation effects, protein was added to the reaction buffer first, and the reaction was initiated by the addition of reporter substrate after 10 minutes of enzyme-compound incubation. The results are reported in Tables 2 and Table 3.

**Table 2.**
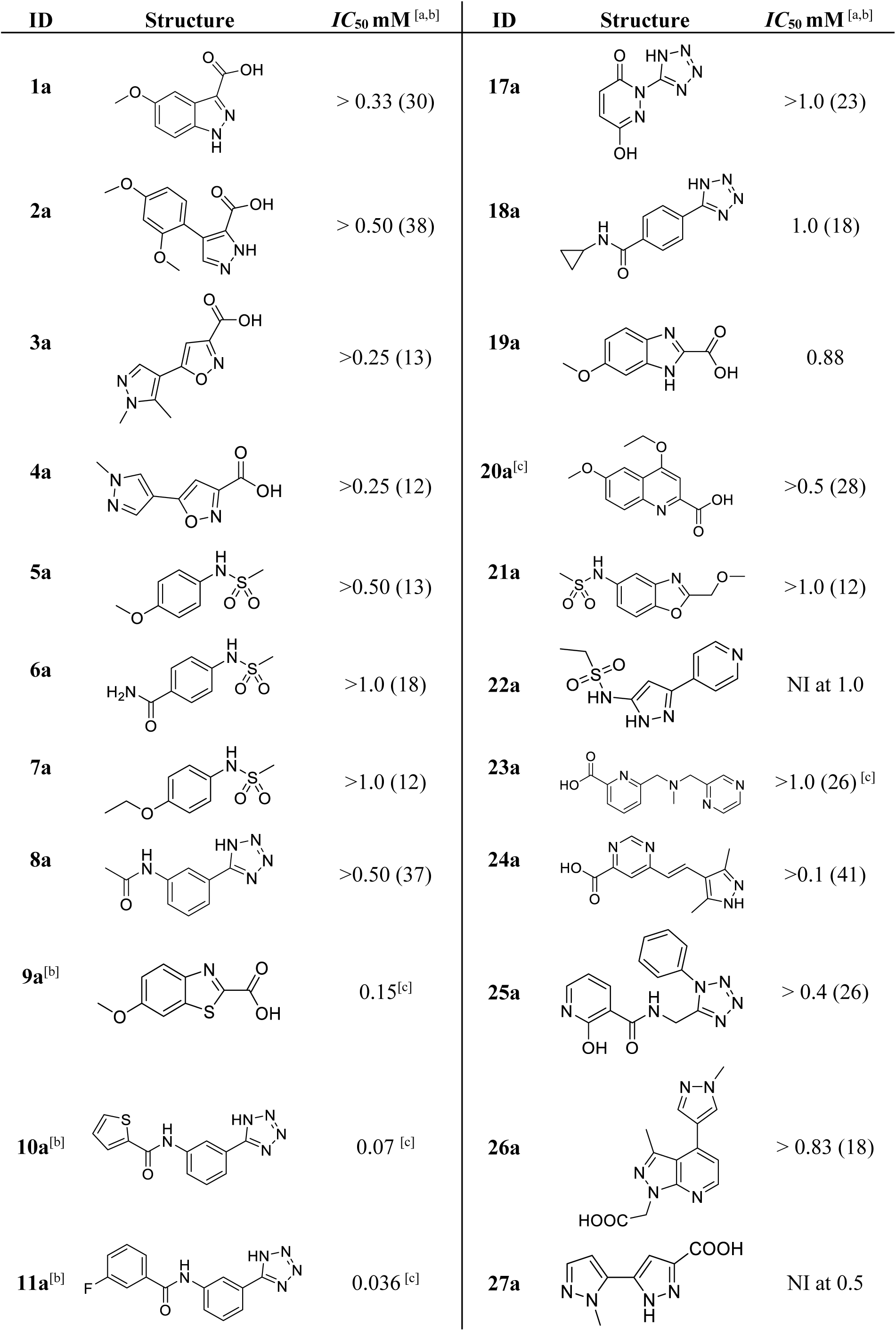
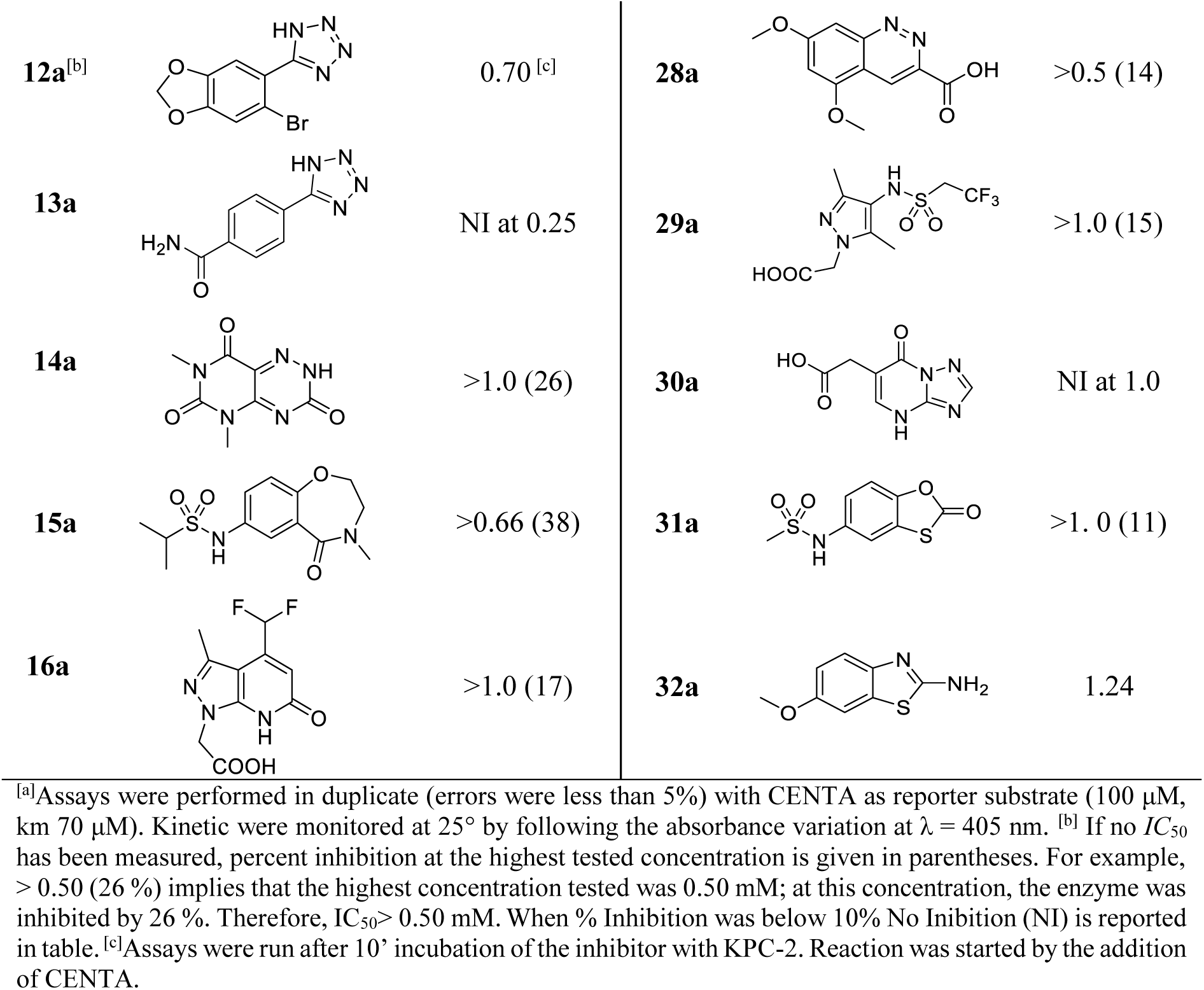
Inhibitory activity of compounds selected from virtual screening.

| ID | Structure | IC<sub>50</sub> mM<sup>[a,b]</sup> | ID | Structure | IC<sub>50</sub> mM<sup>[a,b]</sup> |
| --- | --- | --- | --- | --- | --- |
| 1a |  | > 0.33 (30) | 17a |  | >1.0 (23) |
| 2a |  | > 0.50 (38) | 18a |  | 1.0 (18) |
| 3a |  | >0.25 (13) | 19a |  | 0.88 |
| 4a |  | >0.25 (12) | 20a<sup>[c]</sup> |  | >0.5 (28) |
| 5a |  | >0.50 (13) | 21a |  | >1.0 (12) |
| 6a |  | >1.0 (18) | 22a |  | NI at 1.0 |
| 7a |  | >1.0 (12) | 23a |  | >1.0 (26)<sup>[c]</sup> |
| 8a |  | >0.50 (37) | 24a |  | >0.1 (41) |
| 9a<sup>[b]</sup> |  | 0.15<sup>[c]</sup> | 25a |  | > 0.4 (26) |
| 10a<sup>[b]</sup> |  | 0.07<sup>[c]</sup> | 26a |  | > 0.83 (18) |
| 11a<sup>[b]</sup> |  | 0.036<sup>[c]</sup> | 27a |  | NI at 0.5 |
| **12a**<sup>[b]</sup> |  | 0.70<sup>[c]</sup> | **28a** |  | >0.5 (14) |
| **13a** |  | NI at 0.25 | **29a** |  | >1.0 (15) |
| **14a** |  | >1.0 (26) | **30a** |  | NI at 1.0 |
| **15a** |  | >0.66 (38) | **31a** |  | >1.0 (11) |
| **16a** |  | >1.0 (17) | **32a** |  | 1.24 |

<sup>[a]</sup>Assays were performed in duplicate (errors were less than 5%) with CENTA as reporter substrate (100 μM, km 70 μM). Kinetic were monitored at 25° by following the absorbance variation at $\lambda = 405$ nm. <sup>[b]</sup> If no $IC_{50}$ has been measured, percent inhibition at the highest tested concentration is given in parentheses. For example, > 0.50 (26 %) implies that the highest concentration tested was 0.50 mM; at this concentration, the enzyme was inhibited by 26 %. Therefore, $IC_{50} > 0.50$ mM. When % Inhibition was below 10% No Inhibition (NI) is reported in table. <sup>[c]</sup>Assays were run after 10' incubation of the inhibitor with KPC-2. Reaction was started by the addition of CENTA.

**Table 3:**
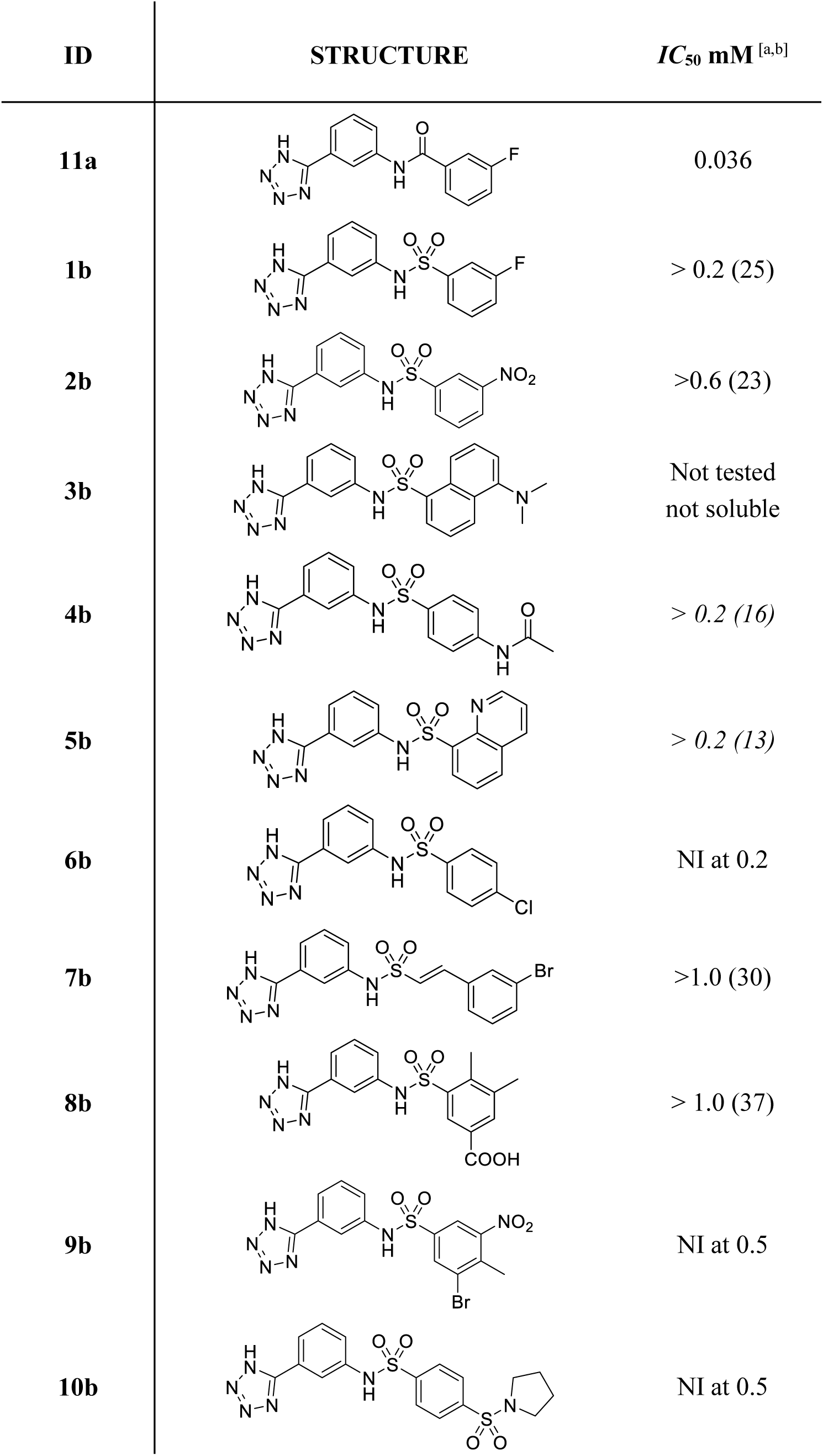
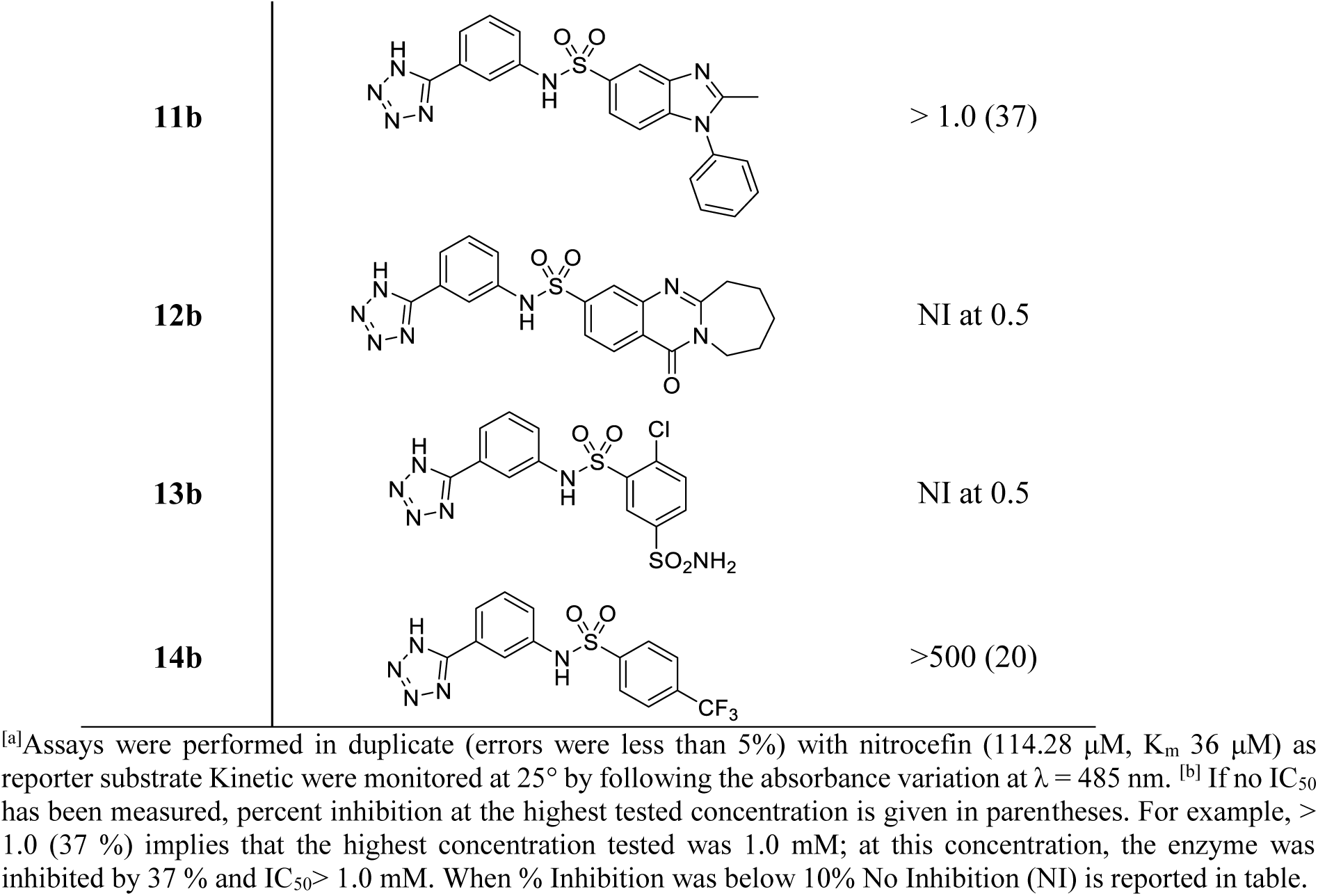
Inhibitory activity of **11a** sulphonamide derivatives.

| ID | STRUCTURE | IC<sub>50</sub> mM<sup>[a,b]</sup> |
| --- | --- | --- |
| 11a |  | 0.036 |
| 1b |  | > 0.2 (25) |
| 2b |  | >0.6 (23) |
| 3b |  | Not tested<br>not soluble |
| 4b |  | > 0.2 (16) |
| 5b |  | > 0.2 (13) |
| 6b |  | NI at 0.2 |
| 7b |  | >1.0 (30) |
| 8b |  | > 1.0 (37) |
| 9b |  | NI at 0.5 |
| 10b |  | NI at 0.5 |
| **11b** |  | > 1.0 (37) |
| **12b** |  | NI at 0.5 |
| **13b** |  | NI at 0.5 |
| **14b** |  | >500 (20) |

<sup>[a]</sup>Assays were performed in duplicate (errors were less than 5%) with nitrocefin (114.28 μM, $K_m$ 36 μM) as reporter substrate. Kinetic were monitored at 25° by following the absorbance variation at $\lambda = 485$ nm. <sup>[b]</sup> If no $IC_{50}$ has been measured, percent inhibition at the highest tested concentration is given in parentheses. For example, > 1.0 (37 %) implies that the highest concentration tested was 1.0 mM; at this concentration, the enzyme was inhibited by 37 % and $IC_{50} > 1.0$ mM. When % Inhibition was below 10% No Inhibition (NI) is reported in table.

Competitive inhibition mechanism and the *K*_i_ for compound **9a** was determined by Lineweaver–Burk (LB) and Dixon plots. For compound **11a**, already reported as competitive inhibitor of the extended spectrum β-lactamase (ESBL) CTX-M15, the *K*_i_ was calculated by the Cheng-Prusoff equation (Ki= IC_50_/(1+ [S]/K_M_) assuming competitive inhibition [36].

## Synthetic procedures

All commercially available chemicals and solvents were reagent grade and were used without further purification unless otherwise specified. Reactions were monitored by thin-layer chromatography on silica gel plates (60F-254, E. Merck) and visualized with UV light, cerium ammonium sulfate or alkaline KMnO_4_ aqueous solution. The following solvents and reagents have been abbreviated: ethyl ether (Et_2_O), dimethyl sulfoxide (DMSO), ethyl acetate (EtOAc), dichloromethane (DCM), methanol (MeOH). All reactions were carried out with standard techniques. NMR spectra were recorded on a Bruker 400 spectrometer with ^1^H at 400.134 MHz and ^13^C at 100.62 MHz. Proton chemical shifts were referenced to the TMS internal standard. Chemical shifts are reported in parts per million (ppm, δ units). Coupling constants are reported in units of Hertz (Hz). Splitting patterns are designed as s, singlet; d, doublet; t, triplet; q quartet; dd, double doublet; m, multiplet; b, broad. Mass spectra were obtained on a 6520 Accurate-Mass Q-TOF LC/MS and 6310A Ion TrapLC-MS(n).

### General procedure for the synthesis of sulfonamides 1-6b

To a solution of 3-(1H-tetrazol-5-yl)aniline (1 eq.) in DCM dry (25 mL) at room temperature and under nitrogen atmosphere, pyridine (3 eq.) and the appropriate sulfonyl-chloride (1.2 eq.) were added. The mixture was reacted at room temperature for 2-12 h. The reaction was quenched with aqueous satured solution of NH_4_Cl and acidified at pH 4 with aqueous 1N HCl. The aqueous phase was extracted with AcOEt, and the organic phase washed with brine, dried over Na_2_SO_4_ and concentrated. The crude was crystalized from MeOH or Et_2_O to give the desired product.

### N-(3-(1H-tetrazol-5-yl)phenyl)-3-fluorobenzenesulfonamide (1b)

Pale yellow solid (150 mg, yield 47%). ^1^H NMR (400 MHz, DMSO-d6) δ 7.19 (dd, *J* = 2.2, 8.1 Hz, 1H), 7.33 – 7.45 (m, 2H), 7.47 – 7.58 (m, 3H), 7.62 (d, *J* = 7.7 Hz, 1H), 7.76 (t, *J* = 1.8 Hz, 1H), 10.61 (s, 1H), the H of tetrazole exchanges. MS m/z [M+H]^+^ 320.1; [M-1]^−^ 318.0.

### N-(3-(1H-tetrazol-5-yl)phenyl)-3-nitrobenzenesulfonamide (2b)

Pink solid (62% yield). ^1^H-NMR (400 MHz, DMSO-d6) δ: 7.31 (ddd, J = 1.0, 2.3, 8.2 Hz, 1H), 7.51 (t, J = 8.0 Hz, 1H), 7.74 (dt, J = 1.2, 7.8 Hz, 1H), 7.82 – 7.96 (m, 2H), 8.18 (dt, J = 1.3, 7.9 Hz, 1H), 8.46 (ddd, J = 1.0, 2.3, 8.2 Hz, 1H), 8.54 (t, J = 2.0 Hz, 1H), 10.90 (s, 1H); the H of tetrazole exchanges. ^13^C-NMR (DMSO-d6) δ: 117.35, 120.09, 123.04, 127.35, 129.17, 129.22, 129.69, 133.21, 136.59, 137.99, 139.81, 148.82, 154.28. MS m/z [M+H]^+^ Calcd for C_13_H_10_N_6_O_4_S: 346.0 Found: 347.2.

### N-(3-(1H-tetrazol-5-yl)phenyl)-5-(dimethylamino)naphthalene-1-sulfonamide (3b)

White solid (31% yield). ^1^H NMR (400 MHz, DMSO-d6) δ 2.78 (s, 6H), 7.20 (dd, J = 1.5, 7.6 Hz, 1H), 7.24 – 7.38 (m, 3H), 7.39 – 7.65 (m, 3H), 8.27 (dd, J = 1.6, 7.5 Hz, 1H), 8.41 (ddd, J = 1.5, 7.5, 19.3 Hz, 2H), the H of tetrazole exchanges. MS m/z [M+H]^+^ Calcd for C_19_H_18_N_6_O_2_S: 394.1 Found: 395.1.

### N-(4-(N-(3-(1H-tetrazol-5-yl)phenyl)sulfamoyl)phenyl)acetamide (4b)

Pink solid (52% yield). ^1^H NMR (400 MHz, DMSO-d6) δ 2.12 (s, 3H), 7.31 (ddd, J = 1.0, 2.3, 8.2 Hz, 1H), 7.44 (t, J = 7.9 Hz, 1H), 7.65 – 7.72 (m, 3H), 7.74 – 7.79 (m, 2H), 7.86 (t, J = 1.9 Hz, 1H), the H of tetrazole exchanges. ^13^C NMR (100 MHz, DMSO-d6) δ 22.58, 118.86, 118.95, 122.60, 123.07, 126.32, 127.96, 128.87, 129.96, 133.69, 139.01, 142.97, 170.57. MS m/z [M+H]^+^ Calcd for C_15_H_14_N_6_O_3_S: 358.1 Found: 359.2.

### N-(3-(1H-tetrazol-5-yl)phenyl)quinoline-8-sulfonamide (5b)

White solid (88% yield). ^1^H NMR (400 MHz, Acetone-d6) δ 7.28 – 7.42 (m, 2H), 7.59 – 7.90 (m, 4H), 8.03 (dt, J = 1.1, 1.8 Hz, 1H), 8.25 (dd, J = 1.5, 8.2 Hz, 1H), 8.42 (dd, J = 1.4, 7.3 Hz, 1H), 8.52 (dd, J = 1.8, 8.4 Hz, 1H), 9.21 (dd, J = 1.8, 4.3 Hz, 1H), 9.41 (s, 1H), the H of tetrazole exchanges. ^13^C NMR (100 MHz, Acetone-d6) δ 117.35, 120.09, 123.01, 123.04, 125.55, 128.06, 129.17, 129.69, 129.79, 130.11, 133.27, 139.81, 140.02, 140.84, 149.72, 154.28. MS m/z [M+H]^+^ Calcd for C_16_H_12_N_6_O_2_S: 352.1 Found: 353.2.

### N-(3-(1H-tetrazol-5-yl)phenyl)-4-chlorobenzenesulfonamide (6b)

Light yellow solid (93% yield). ^1^H NMR (400 MHz, Methanol-d4) δ 7.57 (t, J = 8.0 Hz, 1H), 7.74 (dt, J = 1.3, 7.9 Hz, 2H), 7.78 – 7.84 (m, 1H), 7.99 (d, J = 8.6 Hz, 2H), 8.06 (d, J = 8.6 Hz, 2H), 8.41 (t, J = 1.9 Hz, 1H), the H of tetrazole exchanges. ^13^C NMR (100 MHz, Methanol-d4) δ 117.35, 120.09, 123.01, 123.04, 125.55, 128.06, 129.17, 129.69, 129.79, 130.11, 133.27, 139.81, 140.02, 140.84, 149.72, 154.28. MS m/z [M+H]^+^ Calcd for C_13_H_10_ClN_5_O_2_S: 335.0, 337.0 Found: 336.1, 338.2.

## Results and Discussion

### Virtual Screening

The binding sites in the available KPC-2 crystal structures were analyzed to select a suitable receptor for docking. Alignment and superposition of the binding site residues of the seven available crystal structures of *E.coli* and *K. pneumoniae Kp*KPC-2 revealed a rather rigid binding site with only Trp105 adopting two different rotamers, a closed conformation found 6-times and an open one, found two-times. In one structure, both rotamers were present (Fig. 4). Thus, for virtual screening, the structure with the highest resolution was selected (*K. pneumoniae* KPC-2 in complex with the covalent inhibitor penamsulfone PSR-3-226 (PDB code 3RXW), 1.26 Å resolution). This structure contained both rotamers of Trp105. For virtual screening, the closed conformation was chosen, as this is the most dominant conformation upon ligand binding.

**Figure 4.**
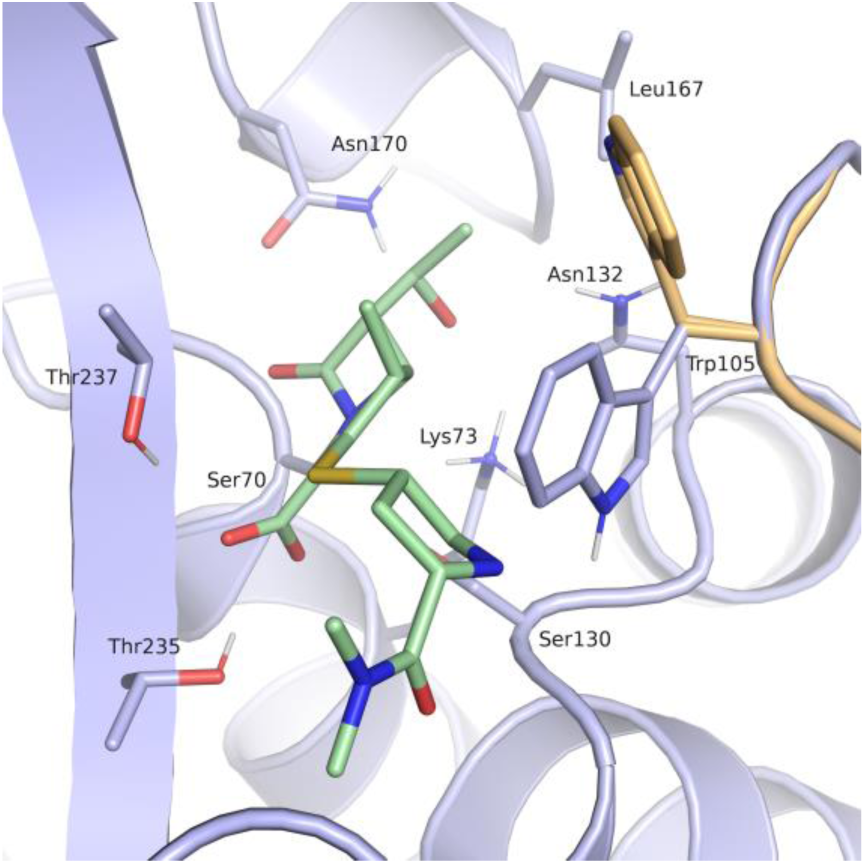
Binding site of *Kp*KPC2 (PDB code 3RXW) with ligand meropenem (green, PDB code 4EUZ). The receptor conformation used for docking is coloured in blue, the rotamer of Trp105 not considered in the docking setup in beige (Ambler numbering) [22].

Only little diversity with respect to bound ligands was found in the *Kp*KPC2 structures. To obtain a more detailed picture on key interactions and to derive a pharmacophore hypothesis, PoSSuM - Search K was used to search for similar binding sites containing non-covalent ligands. This resulted in thirteen structures (Table 1), all having tetrazoles or carboxylates derivatives bound in the hydrophilic pocket formed by the amino acids corresponding to Thr235, Thr237, Ser130 and Ser70 in *Kp*KPC2 (Fig.4). Seven of the contained ligands were fragment hits for *E.coli* CTX-M class A extended spectrum β-lactamase (ESBL), and four were derivatives of the most potent screening hit. Further, a structure of *S. fonticola* SFC-1 S70A β-lactamase in a non-covalent complex with meropenem and one of *E.coli* Toho-1 R274N: R276N β-lactamase in complex with a boronic acid were retrieved. Superposition of the binding site residues of *Kp*KPC-2 (PDB code 3RXW) and the CTX-M β-lactamase structures gave rmsd values for the Cα atoms between 0.72 and 0.82 Å, for superposition of *Kp*KPC-2 and *S. fonticola* SFC-1 (PDB code 4EUZ) 0.28 Å and for superposition *Kp*KPC-2 and *E.coli* Toho-1 (PDB code 4BD0) 0.73 Å (Fig. 5).

**Figure 5.**
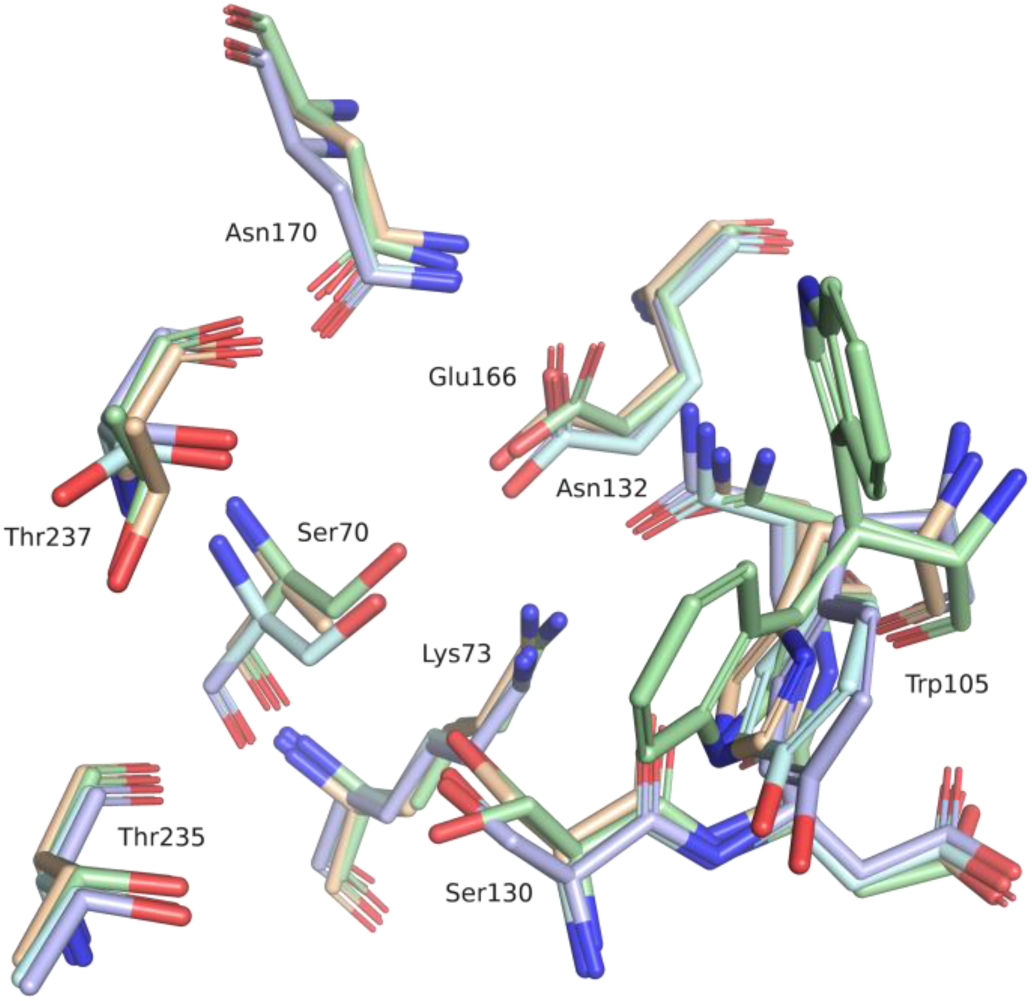
Superposition of the *Kp*KPC-2 binding site residues (PDB code 3RXW, green) with the *E.coli* CTX-M-9 β-lactamase (PDB code 4DDS, blue), the *S.fonticola* SFC-1 β-lactamase (PDB code 4EUZ, beige) and the *E.coli* TOHO-1 β-lactamase (PDB code 4BD0, cyan) (Ambler numbering) [22].

Based on the retrieved structures, a pharmacophore hypothesis was derived. All of the ligands in these structures as well as the β-lactamase binding protein (PDB code 3E2L, 3E2K) and the covalent ligand of the structure used as receptor, formed a hydrogen-bond with Thr235 or Thr237. Accordingly, a hydrogen-bond acceptor at the corresponding ligand position was considered to be crucial for binding (Fig. 3). Further, in most of the structures the ligands formed interactions with Trp105 (*Ambler* numbering) [22]. Therefore, this interaction was also included in the pharmacophore hypothesis. Hydrogen-bond interactions to Asn130 were found in four structures (PDB codes 3RXW, 3G32, 3G30, 4EUZ) and included as well.

A hierarchical approach was adopted for virtual screening. First, our in-house database of around five million purchasable compounds was filtered for lead-like molecules [23]. In the second step, the obtained hits were screened with the above-described pharmacophore resulting in 44658 compounds. Out of these, 31122 compounds could be docked into the *Kp* KPC2 binding site. Filtering these binding poses again with the pharmacophore resulted in 2894 compounds. These were divided into three clusters, depending on the functional group placed in the hydrophilic pocket (tetrazoles, carboxylates, sulfonamides) and inspected by eye. Finally, 31 compounds were selected for hit validation (Table 2). Most of the selected chemotypes carried an anionic group, mainly a carboxylic group or its bioisostere, the tetrazole ring. Candidates were predicted to orient the anionic side of their moiety in the carboxylic acid binding site of KPC-2, delimited by Ser130, Thr235 and Thr237 and present in all serine-based Beta-lactamases. In the above mentioned site, in fact, binds the C(3)4’ carboxylate of β-lactams antibiotics as well as the sulfate group of avibactam and the carboxylic group of other known BLs inhibitors [12,32,37,38,39].

### Hit Evaluation

The majority of the selected candidates were fragment-like as defined by the “rule of three” [14]. Thus, potencies in the high micromolar to millimolar range were expected. Unfortunately, the required high concentrations for ligand testing could not always be achieved due to solubility issues which might have resulted false negatives after testing. However, some of the tested molecules inhibited the hydrolytic activity of KPC-2 with millimolar potency. Among those, compounds **9a** and **11a** were the most promising compounds with micromolar affinities (*IC*_*50*_ of 0.15 and 0.036 mM, translating to ligand efficiencies (LE) of 0.38 and 0.28 kcal/mol/non-hydrogen atom, respectively; Table 2) and were thus further investigated.

Compound **9a** was predicted to place its carboxylate group in proximity of the catalytic Ser70, in the carboxylic acid binding site mentioned above, forming hydrogen bond interactions with the side chains corresponding to amino acids Ser130, Thr235 and Thr237 (Fig. 6). Thr 237 in KPC-2 is known to be necessary for cephalosporinase and carbapenemase activity and is involved in clavulanic acid, sulbactam and tazobactam recognition.[7] This position in β-lactamases that do not have carbapenemase or extended-spectrum b-lactamase (ESBL) activity generally corresponds to an alanine. The side chain hydroxyl groups of Ser130 and Ser70 were predicted to form interactions with the nitrogen of the benzothiazole ring. The predicted position of the aromatic system is well placed to establish ring-ring interactions with Trp105, a residue involved, in turn, in the stabilization of β-lactams through mainly hydrophobic and van der Waals interactions (centroids distances of 4.4 and 4.5 A between Trp105 and the thiophene and the benzene rings respectively) The role of Trp105 in substrate and inhibitor interactions in KPC-2 β-lactamase has been deeply investigated being essential for hydrolysis of substrates.[7] The methoxy group of the molecule is oriented towards a rather open and solvent accessible area of the binging site and does not contact any of the surrounding residues. Interestingly, the presence of the sulphur atom of the benzothiazole system seems critical for affinity as the related compound **19a**, the benzimidazole analog, resulted 6-fold less active. Similar, the presence of the carboxylic group appeared to be crucial as compound **32a,** without such a functionality, was 8-fold less active.

**Figure 6.**
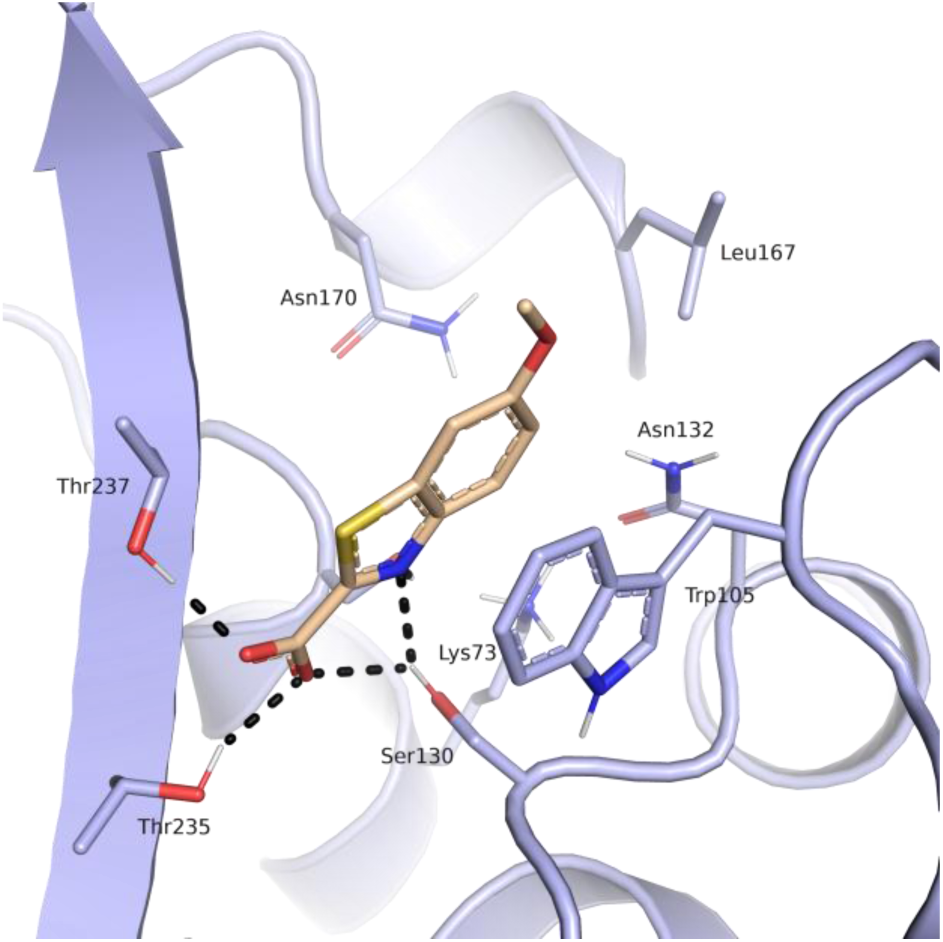
Predicted binding mode of compound **9a** (beige) in the *Kp*KPC-2 receptor (blue). Putative hydrogen bond interactions are indicated as black dots (Ambler numbering)[22].

For compound **9a** binding affinity and mode of inhibition was determined by using gradient concentrations of CENTA. Fitting of the obtained data showed that compound **9a** behaves as a competitive inhibitor with a determined *Ki* of 112.0 µM (Fig. 7). Its binding affinity was also determined towards other class A β-lactamases (*IC*_*50*_ vs CTX-M9 160 µM). For this compound aggregating behavior was also excluded by dynamic light scattering experiment (data not shown) [40]. Compound **9a** with its fragment-like characteristic (MW 208.21, determined *Ki* 112.0 µM, LE 0.38 kcal/mol/non-hydrogen atom) exerts an interesting activity *vs* KPC2-2 and represents a very promising molecule to be directed to hit to lead optimization.

**Figure 7.**
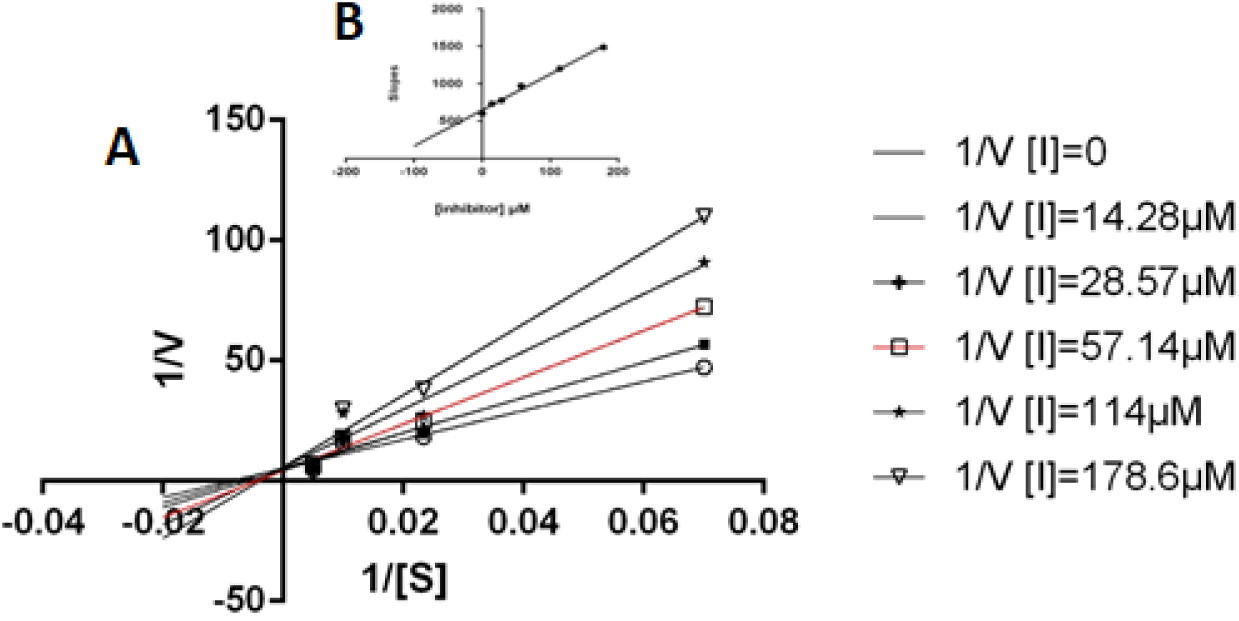
(A) Lineweaver–Burk plot **(A)** and Dixon slope plot **(B)** for competitive inhibitor, compound **9a**.

Among the 32 selected hits evaluated *in vitro* for their binding affinity vs KPC-2, compound **11a** was the most active inhibitor with a micromolar affinity vs KPC-2 (determined *IC*_*50*_ 36 µM, calculated K*i* 14.8 µM, LE 0.28 kcal/mol/non-hydrogen atom).[36] The tetrazole ring of compound **11a**, a well-known bioisostere of the carboxylic group, was predicted to lie in the hydrophilic pocket formed by Thr235, Thr237, Ser130 and Ser70, driving the binding of the inhibitor in KPC-2 active site(Fig. 8). The phenyl ring attached to the tetrazole was predicted to be sandwiched between the Trp105 side with a distance compatible with weak hydrophobic interactions and the backbone of Thr237. The amide group of **11a** was oriented in the canonical site delimited by Asn132, Asn170 and in a further distance Glu166 where the R1 amide side chain of β-lactams is known to bind. However, the amine linker and the second phenyl ring in **11a** were not predicted to form any specific interactions with the protein, except for the amide nitrogen contacting the backbone of Thr237. The distal fluoro-benzene ring was oriented at the entrance of the active site against two hydrophobic patches, one defined by Leu167, closer, and the other by the backbone of Asn170, a residue critical for carbapenemase activity.

**Figure 8.**
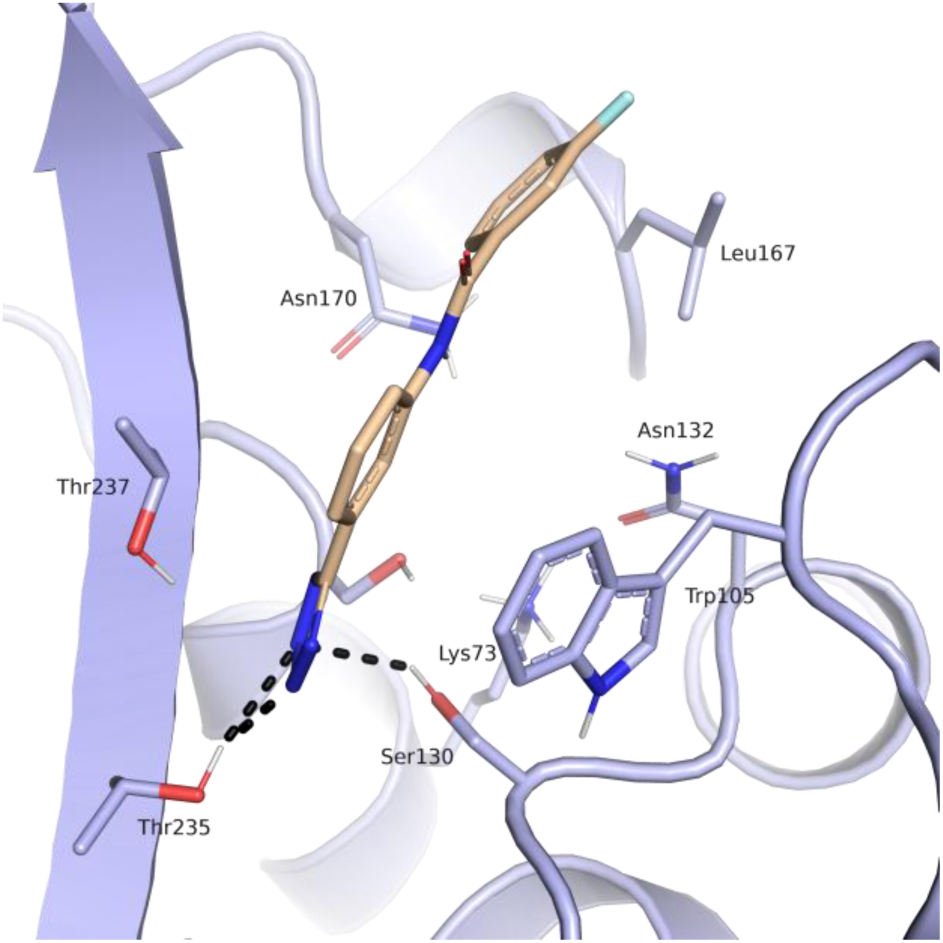
Predicted binding mode of compound **11a** (beige) in the *Kp*KPC-2 receptor (blue). Putative hydrogen bond interactions are indicated as black dots (Ambler numbering)[22].

Based on the predicted binding mode, the tetrazole group of **11a** seemed to be crucial for affinity. (Table 1). Moreover, while the proximal ring appeared to be involved in specific interactions, the amide group and the distal ring did not contact efficaciously the protein. Based on predicted binding mode, chemical size, synthetic accessibility for a rapid structural optimization and ligand efficacy compound **11a** was directed to chemical synthesis development to improve its affinity and to investigate target binding requirements for optimal inhibitor-enzyme interaction.

### Hit derivatization and evaluation

In order to improve the binding affinity of **11a,** the compound was subjected to a hit optimization program. Therefore, the phenyl-tetrazole moiety, that seemed to strongly drive the binding, was retained unaltered, whereas structural modifications on the linker and on the distal aromatic ring were introduced in order to explore and maximize the interactions with the pocket formed by Asn132, Asn170 and Leu167 (Fig. 6). Because the amide linker does not contact efficaciously the protein we chose to replace it with a sulfonamide (Fig. 9). We meant to target residues proximal to the opening of the active site while investigating the potentiality for sulfonamide derivatives.

**Figure 9.**
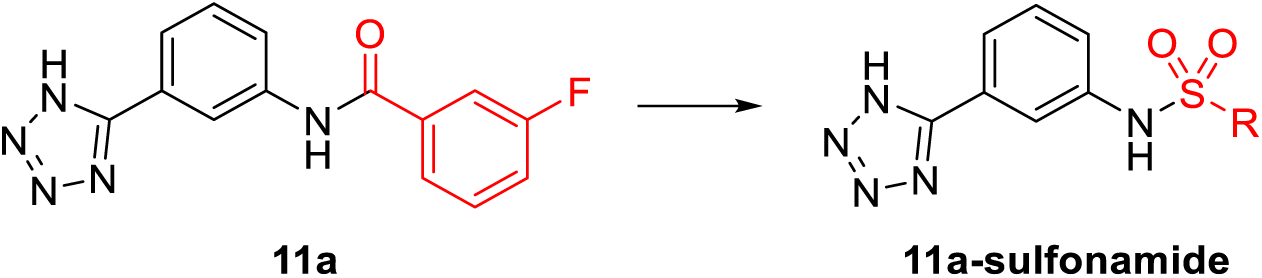
Virtual Screening hit **11a** (left), amine 1 (black) and the optimized part of the molecule (red).

Sulfonamides are more stable towards hydrolysis than carboxyamides, possess an additional hydrogen bonding oxygen atom and their NH is a strong hydrogen bond donor. In addition, the dihedral angle ‘ω’ OSNH measures around 90° compared with the 180° ‘ω’ OCNH angle of amide. Sulfonamides, in addition, have a non-planar configuration that could orient the distal ring towards Leu167 and Asn170 (Fig. 10). Therefore, the introduction of a sp^3^ geometry could allow a more efficacious spanning of the active site compared to the planar amide [41]. Moreover, modeling suggested that the sulfonamide group could form an additional hydrogen bond with Asn132 residue actively involved in substrate recognition and hydrolysis.

**Figure 10.**
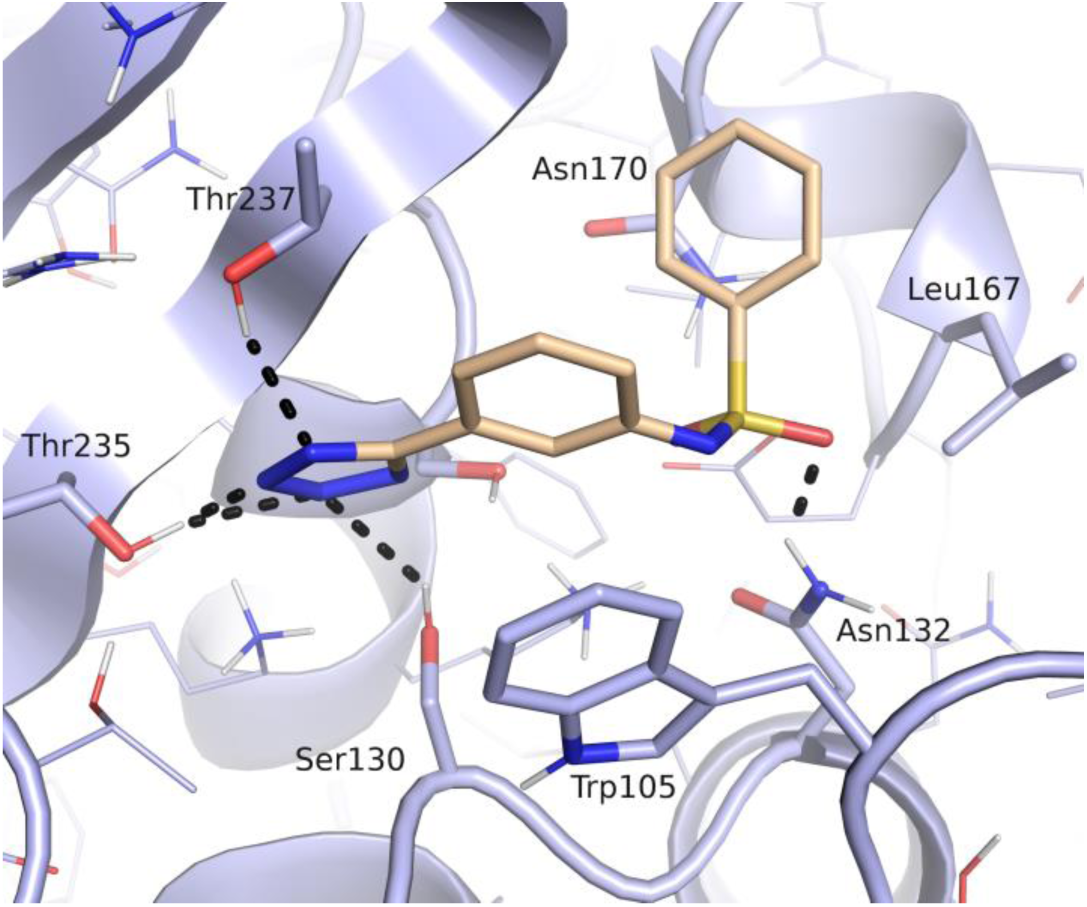
Predicted binding mode of a sulfonamide derivative of compound **11a** (beige) in the *Kp*KPC-2 receptor (blue). Putative hydrogen bond interactions are indicated as black dots (Ambler numbering)[22].

Further, we explored different substitutions on the sulfonamide linker to probe binding interactions. Substituents with different electronic and steric properties (i.e. halogens, nitro, sulfonamide, carboxylic acid, methyl, acetamide, amino groups) were inserted in the different position of the aromatic ring. In addition, the benzene ring was replaced by heterocyclic or extended benzofused systems such as benzimidazole, quinazolinone, naphthalene, or quinolone ring. Based on the availability of compound or building blocks, 6 compounds (**1b-6b**) were synthesized and 8 compounds (**7b-14b**) were purchased to test our hypothesis (Table 3). The fourteen new compounds were tested *in vivo vs* clinical strains overproducing KPC-2 to evaluate their ability to restore bacteria susceptibility to carbapenem meropenem (Table 4).

**Table 4:**
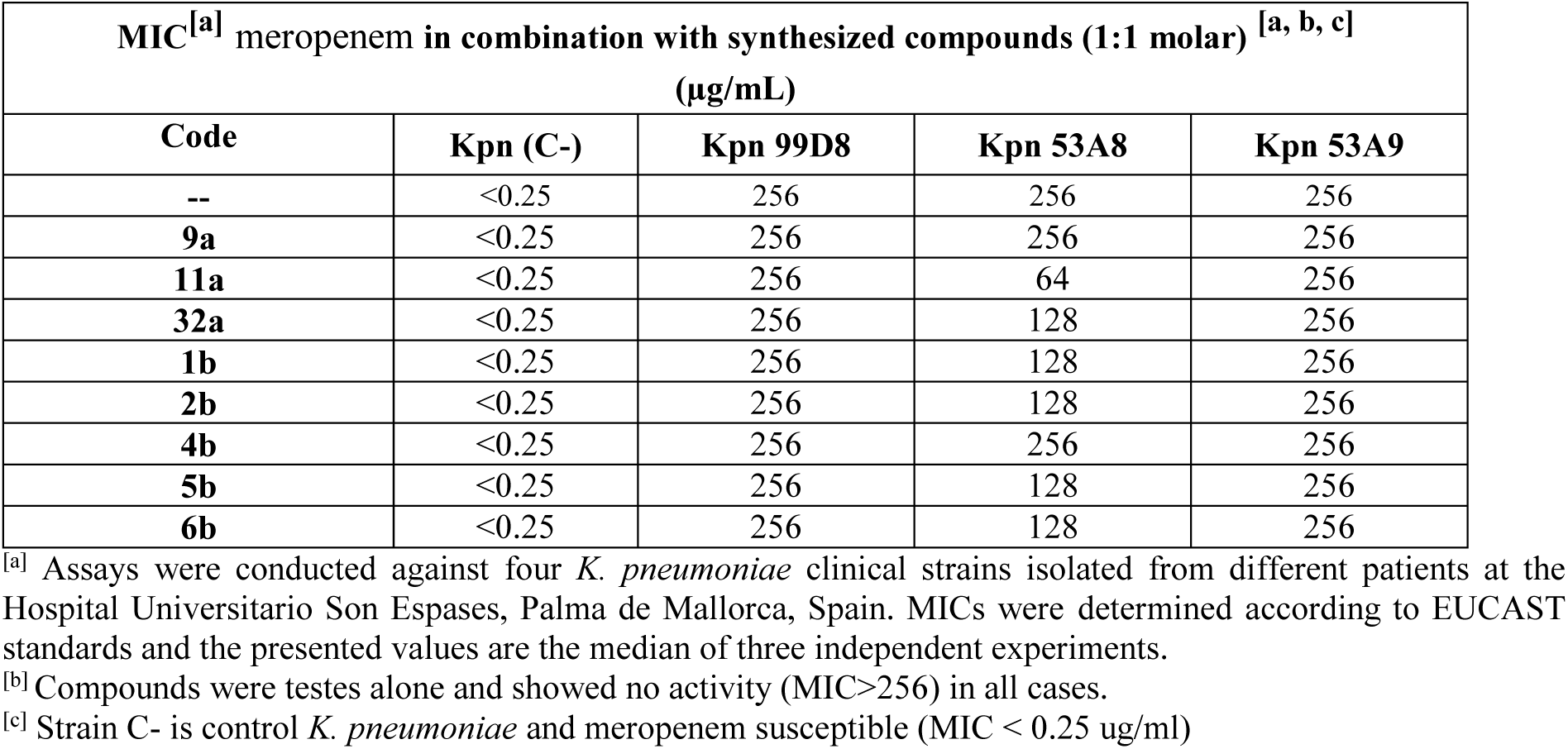
*In vitro* interaction between meropenem and synthesized compounds vs *K. pneumoniae* clinical strains.

Compounds **1b-6b** were synthesized in high yield (75-95% yield) and purity (>95%) through direct reaction of 3-(1H-tetrazol-5-yl) aniline and the appropriate sulfonyl chloride in dichloromethane at room temperature for 3 hours (Fig. 11).

**Figure 11.**
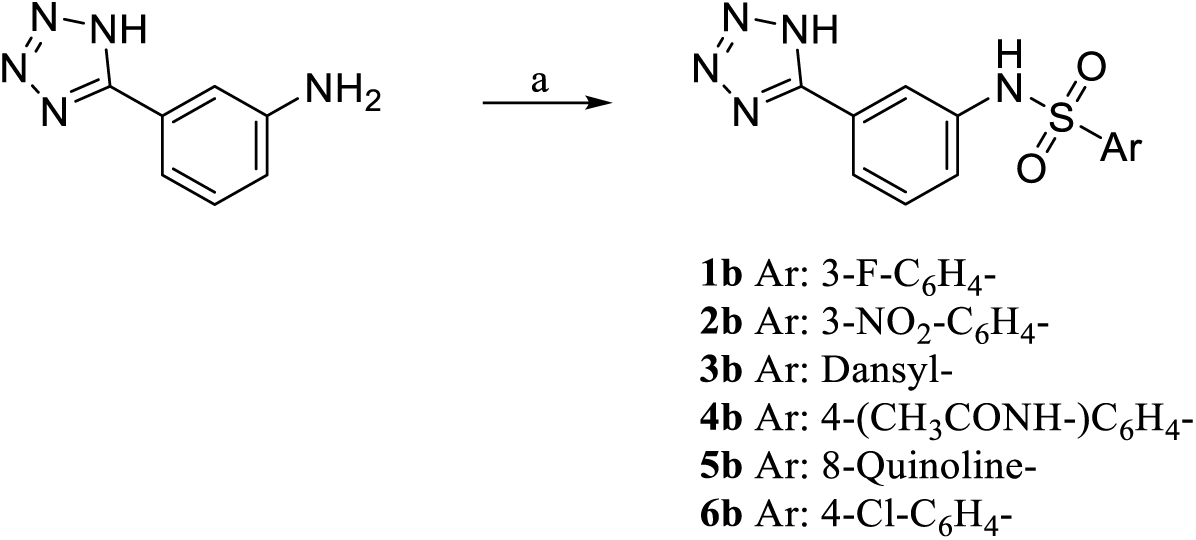
Reagents and conditions. a) aryl-sulfonyl chloride (1.2 eq.), pyridine (3 eq.), dry DCM, N2, r.t, 3 h, 75-95% yield.

The derivatives of compound **11a** were tested for KPC-2 affinity (Table 3). They exhibited either weaker activities than **11a** or were not active at all at tested concentration. Thus, it appeared that the sulfonamide linker is not a suitable group to optimize the affinity of this compound series.

The antimicrobial activity of the best hits **9a** and **11a** and their derivatives was studied in bacterial cell cultures to investigate their ability to cross the outer membrane reaching the periplasmic space, where KPC-2 is secreted and confined in Gram negative bacteria. Compounds were tested for synergy with the β-lactam antibiotic meropenem against four *K. pneumoniae* clinical strains, isolated from different patients at the Hospital Universitario Son Espases, Palma de Mallorca, Spain. One of the four clinical strains was not a KPC-2 producer and was susceptible to meropenem (strain **Kpn (C-)**; MIC <0.25 ug/mL). The three additional strains harbored the blaKPC-2 gene and were resistant to meropenem (Table 4). [32] Noteworthy none of the tested compounds had intrinsic antibiotic activity (MIC >256 ug/mL), against the employed strains, included the susceptible one. The results show that in most cases the compounds were not able to reverse antibiotic resistance and did not showed synergism with meropenem. However, against strain Kpn 53A8 the MIC value was lowered by a factor of two when meropenem was combined with compounds **32a**, **1b**, **2b**, **5b** and **6b** while in combination with compound **11a** the MIC value was reduced by 4 fold.

## Conclusions

In this study two novel KPC-2 inhibitors were identified via an *in-silico* approach. 14 novel tetrazole derivatives originating from the best, low micromolar, hit **11a** were designed, synthesized and evaluated for their ability to inhibit KPC-2. We introduced chemical diversity on the distal part of the inhibitor, choosing to keep unchanged the anchor tetrazole ring while modifying the amide and the distal ring. The results suggests that a sulfonamide linker is not suitable to improve the potency of **11a**. Future optimization work should instead rather concentrate on exploring secondary binding sites more distal from the pocket that the screening hits are supposed to address [42]. If the amide functionally found in **11a** or alternative linkers are best suited remains to be explored. Nevertheless, two promising hit compounds for KPC-2 were retrieved which can serve as starting points to derive more potent inhibitors.

Although a decrease of potency in *in vitro* tests was registered for the designed and synthesized chemical entities, as none of the compounds was able to trigger stronger interactions with the open region of KPC-2 they were meant to target, this study yielded a better comprehension of the catalytic pocket of this enzyme. Our study provided a better understanding of how challenging the target of additional, superficial, binding pockets is and how it could be critical in designing inhibitors with improved potency, especially in area proximal to the active site opening.

The (1H-tetrazol-5-yl)phenyl ring was most frequents among the high scoring candidates in our *in silico* study, suggesting that this functionality is well suited to anchor ligands in the in KPC-2 binding site. The rather weak affinity of the ligands hints that rest of the ligand, i.e. the functional groups out of the center phenyl ring and the amide liker, need to be optimized to increase potency. However, introducing a sulfonamide linker was detrimental for potency. We hypothesize that the presence of a sulfonamide instead of an amide led to a rearrangement of the ligand in the binding site to minimize steric hindrance, and thus resulted in the loss of key interactions. These rearrangements can be particularly critical in non-covalent inhibitors like ours that are not stabilized by a covalent interaction with the catalytic serine, as this type of inhibitors are supposed to have lower residence times with respect to covalent β-lactamase inhibitors (Fig.1). In designing larger and more potent inhibitors, additional secondary binding sites which have been found to be critical for affinity improvement need to be considered [42]. Further medicinal chemistry work is ongoing to significantly increase the potency of the most promising compound **11a** *in vitro* and *in vivo* and new chemistry is under evaluation for these derivatives, taking advantage of other additional recognition sites in KPC-2.

## Acknowledgements

We thank Hayarpi Torosyan from Brian K Shoichet’s Laboratory at USCF for dynamic light scattering measurements on compound **9a**, Josef Kehrein for contributions to the modelling work, and Openeye for free software licenses. Virtual screening was performed on resources provided by UNINETT Sigma2 - the National Infrastructure for High Performance Computing and Data Storage in Norway and the Mogon cluster of the Johannes Gutenberg University Mainz.

